# A diverse repertoire of anti-defense systems is encoded in the leading region of plasmids

**DOI:** 10.1101/2023.02.15.528439

**Authors:** Bruria Samuel, David Burstein

## Abstract

Plasmids are an important source of antibiotic-resistance genes that mobilize horizontally between bacteria, including many human pathogens. Bacteria express various defense mechanisms, such as CRISPR-Cas, restriction-modification systems, and SOS-response genes, to prevent the invasion of mobile elements. Yet, plasmids efficiently and robustly overcome these defenses during conjugation. Here, we show that the leading region of plasmids, which is the first to enter recipient cells, is a hotspot for an extensive repertoire of anti-defense systems, encoding anti-CRISPR, anti-restriction, anti-SOS, and other counter-defense proteins. We further demonstrate that focusing on these specific functional regions can lead to the discovery of diverse anti-defense genes. Promoters known to allow expression from ssDNA were prevalent in the leading regions, potentially facilitating rapid protection against bacterial immunity in the early stages of plasmid invasion. These findings reveal a new facet of plasmid dissemination and provide theoretical foundations for developing conjugative delivery systems for natural microbial communities.

## Introduction

Bacterial conjugation is a major horizontal gene transfer (HGT) mechanism in which DNA is transferred from a donor to a recipient cell by direct contact. Conjugative plasmids are an important driver of rapid bacterial evolution^1,2^, and they represent one of the gravest challenges for counteracting antimicrobial resistance genes (ARGs) dissemination^3,4^.

The DNA transport machinery of conjugative elements includes type IV secretion system (T4SS) proteins, an origin of transfer (*oriT*), and a DNA-processing nucleoprotein complex called the relaxosome, composed of the relaxase and often additional auxiliary proteins necessary for conjugation. While conjugative plasmids and integrative conjugative elements (ICEs) encode for the entire transport machinery, some mobile elements contain only the relaxase gene and an *oriT* sequence but lack the machinery required for self-transfer^5^. Such elements, termed “mobilizable plasmids”, can be transferred by exploiting the conjugation machinery of a co-residing conjugative element^6,7^. We will refer to both conjugative and mobilizable elements as “potential conjugative elements”, as they all have the potential to be transferred to another cell by conjugation^8^.

Conjugation initiation occurs when the relaxosome is assembled on the plasmid’s *oriT*, and the relaxase nicks the *nic* site, located within the *oriT*^9,10^. The nicked DNA strand is then transferred with the covalently attached relaxase through the T4SS into the recipient cell. The region first transferred to the recipient cell is defined as the leading region. The relaxase gene is typically located on the plasmid in close proximity to the *oriT*, on the lagging region, which is the last to enter the recipient cell^11,12^.

Previous studies suggested that genes in the leading region are important for plasmid stability during conjugation^13,14^. It was demonstrated that, in certain conjugative plasmids, leading region genes are expressed early upon the entry of the plasmid into the recipient cell^15–17^, even before the transfer is complete^18^. The leading regions of these plasmids contain promoters designated F*rpo* (*ssiD* and *ssiE* in F plasmid^19^, and *ssi2* and *ssi3* in the IncI1 plasmid Col1b-9^15^). These promoters can adopt a stem-loop structure such that the regions of the −10 and −35 elements mimic a double-strand conformation, allowing their recognition by the host RNA polymerase^15,20^. It was therefore proposed that F*rpo* may serve as a single-stranded promotor that allows early expression of the leading region genes^16,21,22^. Indeed, the transcription levels and gene product accumulation indicate rapid expression of genes located in the leading region^16,23,24^.

Similarly to other foreign genetic material, conjugative elements face a wide repertoire of prokaryotic defense systems, including restriction-modification (R-M), CRISPR-Cas, and more^25,26^. Despite these defense systems, designed to prevent the entry of exogenous DNA, which can impose a fitness cost to the host^27,28^, HGT widely persists across species^29^. This is enabled, inter alia, owing to different mechanisms mobile genetic elements (MGEs) have developed to overcome prokaryotic defense systems, including anti-defense proteins such as anti-restriction and anti-CRISPR proteins^30,31^. Such anti-defense genes were also identified on multidrug-resistance plasmids and were shown to affect their horizontal transfer^32,33^. Interestingly, a few of the genes described in the leading region of plasmids encode proteins with anti-defense functions and are known to be early expressed. These include ArdA, an anti-restriction protein that inactivates type I R-M systems in recipient cells^34^, and PsiB, which inhibits the bacterial SOS stress response during conjugation^24^. However, these studies were performed on very few plasmids from specific types (namely, IncI, ColIb-P9, and F plasmids). Little is known about the majority of the genes in the leading region of conjugative elements and their function during conjugation^19,22^. We hypothesized that the leading region of conjugative elements plays a key role in their ability to cope with defense systems through genes encoding for various anti-defense mechanisms. For these anti-defense genes to be effective, they probably need to be rapidly expressed in the conjugation establishment phase, i.e., in the very early stages of the newly acquired mobile element in the recipient cell, similar to anti-CRISPRs and the *ocr* anti-restriction gene in phage, which are expressed early upon infection^35–39^. Exploring this hypothesis, we discovered that the leading regions are indeed highly enriched with anti-defense genes, and found that these regions contain a plethora of uncharacterized genes, many of which are likely to have yet undescribed anti-defense-related functions. Our results suggest that leading regions of conjugative elements act as “anti-defense islands”, protecting conjugative elements from host defense systems upon entry to the recipient cell.

## Results

### The leading regions of conjugative elements are enriched with anti-defense genes

We aimed to explore the position of anti-defense genes in relation to the origin of transfer (*oriT*). To this end, we analyzed all sequences annotated as plasmids in the NCBI’s Whole Genome Shotgun (WGS) database. Notably, these datasets are enriched with pathogens and other clinically relevant bacteria. We focused our analysis on plasmid sequences containing an *oriT* close to a relaxase/*traM* relaxosome gene, which allowed us to discern between the leading and lagging regions. Relaxosome components are typically encoded in the lagging region adjacent to the *oriT*.^11,12,40^. To reduce the redundancy of plasmids in the databases, we removed highly similar plasmids (see Methods), decreasing the number of plasmids from 3,192 to 1,554 representative sequences (Fig. 1).

**Figure 1.**
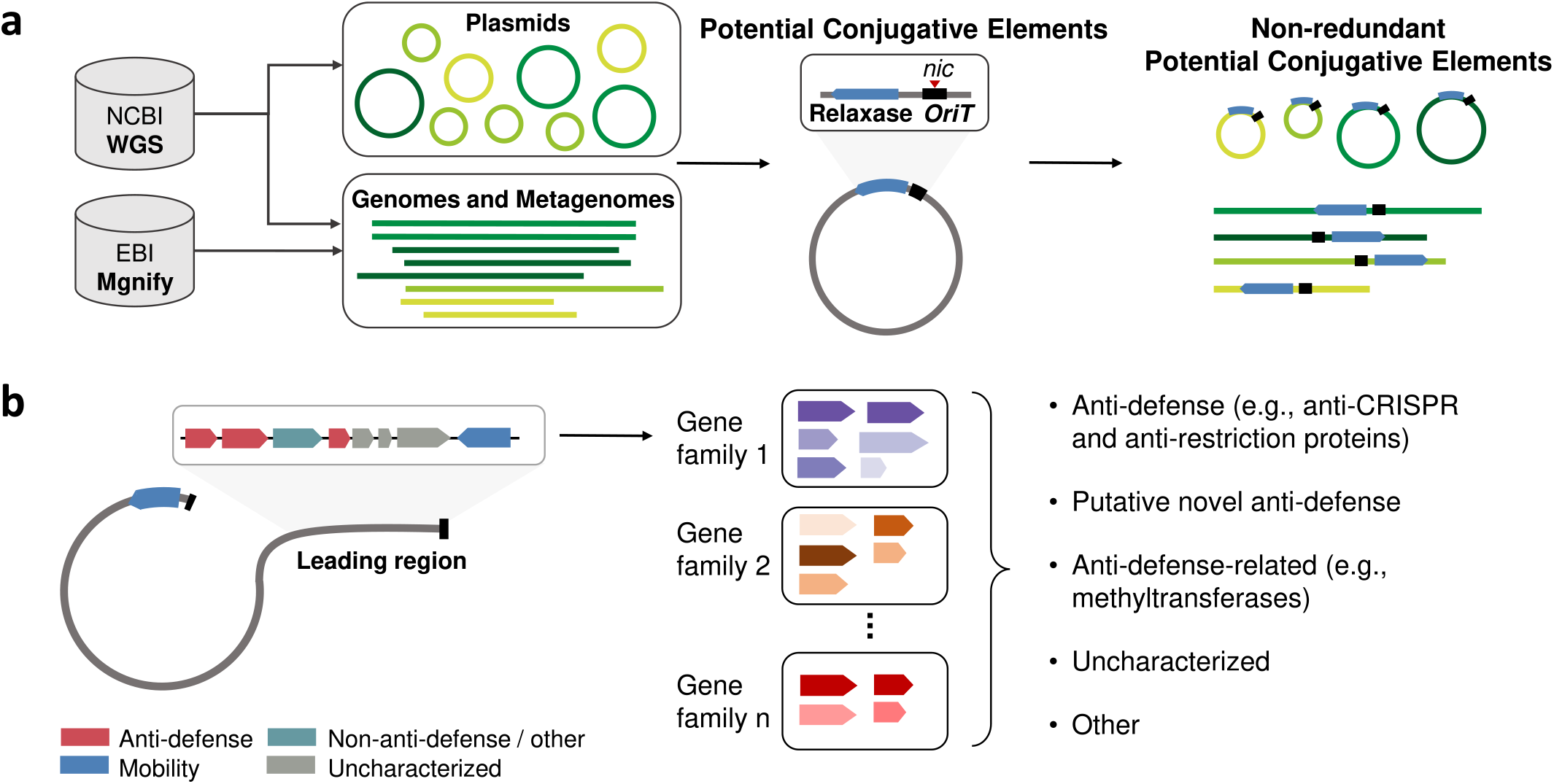
Workflow overview. **(a)** All assembled genomes and metagenomes available in NCBI’s^41^ and EBI’s^42^ databases were analyzed. In the first phase, we considered only sequences explicitly annotated as plasmids. In the second phase, we also included “potential conjugative elements”, i.e., all sequences that contained a detectable relaxosome component gene in proximity to a known *oriT* sequence. We then omitted redundant elements based on sequence similarity. **(b)** We identified the leading and lagging regions of the potential conjugative elements, which enter the recipient cell first and last accordingly, and mapped known anti-defense (e.g., anti-restriction and anti-CRISPR) and transfer-related genes to these sequences. We then focused on the genes encoded in the leading region and characterized various gene families enriched in these regions (anti-defense, putative novel anti-defense, anti-defense-related, uncharacterized genes, and other gene families).

We next searched for known anti-defense genes within these plasmids. These included genes encoding for anti-CRISPR proteins, which antagonize the activity of CRISPR-Cas systems^30^; anti-restriction proteins, inhibiting restriction endonucleases^31,43^; and SOS-inhibitors, which suppress the potentially deleterious host SOS-response elicited by plasmid entry. Notably, the SOS response may also induce the production of nucleases that could provoke the degradation or mutation of the transferred DNA^44–46^.

We measured the relative abundance of anti-defense genes at each position of the plasmid sequences with respect to the location of the *oriT* sequence. The frequency of anti-defense genes at each position revealed that the leading region of these elements is highly enriched with anti-defense genes (Fig. 2a). Specifically, the first 30 ORFs of the leading region were significantly enriched with anti-defense genes (p-value < 0.05; see Methods). Separate analyses performed for each of the anti-defense categories (i.e., anti-CRISPRs, anti-restriction, and SOS-inhibition) showed a similar trend (Fig 2b), and most positions within the first 30 ORFs were significantly enriched (p-value < 0.05; see Methods) in each category.

**Figure 2.**
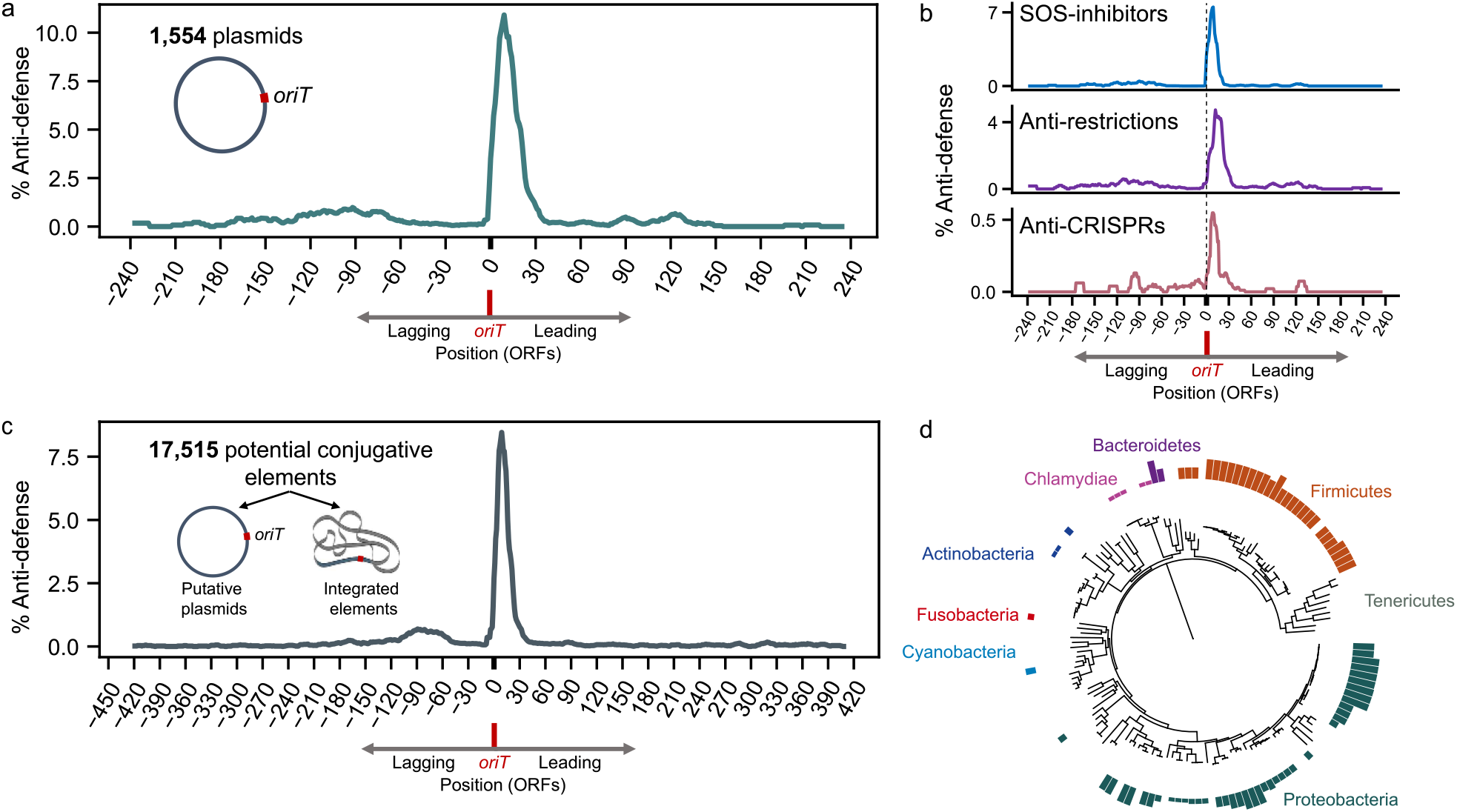
Enrichment of anti-defense genes in the leading regions. **(a)** Analysis of anti-defense proteins encoded on annotated plasmids. The ORF indexes on the *x*-axis are according to their position relative to the origin of transfer (*oriT*), such that 0 is the first ORF in the leading region. Presented are positions that were represented in at least 100 plasmids (Supplementary Fig. 1 include all positions). The *y*-axis denotes the average frequency of anti-defense genes (combining well-characterized SOS inhibitor, anti-restriction, and anti-CRISPR genes) over a window of five ORFs upstream and downstream. **(b)** Breakdown of the anti-defense gene frequency according to their functional categories. **(c)** Analysis of anti-defense gene frequency from potential conjugative elements retrieved from genomic and metagenomic databases. The presented positions are represented in at least 500 sequences (see also Supplementary Fig. 1). **(d)** The host phylogenetic distribution of 13,882 non-redundant plasmids and potential conjugative elements we analyzed. The phylogeny does not include 4,613 elements originating from metagenomes and sequences that could not be mapped to the tree. The plasmid hosts are mapped to the bacterial tree of life^47^, with bars color-coded according to phyla, representing the conjugative element occurrences on a log scale

To test the generality of our findings, we wished to expand our dataset beyond NCBI WGS sequences that were explicitly annotated as plasmids, as these consisted mainly of isolates of pathogens and model organisms and did not include ICEs. We sought potential conjugative elements, which may be unannotated plasmids or ICE-containing sequences, within all publicly available assemblies from NCBI and EBI, including both genomes and metagenomes. We searched in this extensive dataset for sequences with relaxase/*traM* genes in proximity to an *oriT* sequence. After excluding the well-characterized plasmids that we have already analyzed, we found 17,515 additional non-redundant potential conjugative elements. We scanned this set for anti-defense genes and found a very strong enrichment of anti-defense genes encoded in the leading regions of these potential conjugative elements (Fig. 2c). This corroborates our initial findings and demonstrates that, across a large variety of conjugative elements from a wide diversity of bacterial hosts (Fig. 2d; Supplementary Fig. 2), the genes that are transferred first are disproportionally enriched with genes inhibiting the host’s defense systems.

### Uncovering various anti-defense-related genes in the leading region

Given our results, we wondered what could be learned about the function of the most prevalent gene families in the leading regions that were not included in the three anti-defense categories we analyzed. We first clustered the first 30 genes of all 18,489 non-redundant conjugative elements to gene families. Then, to discern between genes that are generally abundant in conjugative elements and genes that are specifically enriched in the leading region, we tested which of the most prevalent gene families within the leading region are significantly enriched in the first 30 ORFs. This analysis revealed that out of the 105 largest families (with more than 450 genes each), 91 were significantly enriched in the first 30 ORFs (Fisher’s exact test; p-value < 0.001 after accounting for multiple testing, see Supplementary Dataset 1).

These 91 gene families included, in addition to known anti-defense genes, also three main anti-defense-related functional groups (Fig. 3a). One of the most prominent functional groups was “orphan” DNA-methyltransferases, which presumably methylate conjugative elements to protect them from the host R-M systems, as previously demonstrated for bacteriophages and proposed for other mobile elements, including plasmids^21,48–51^.

**Figure 3.**
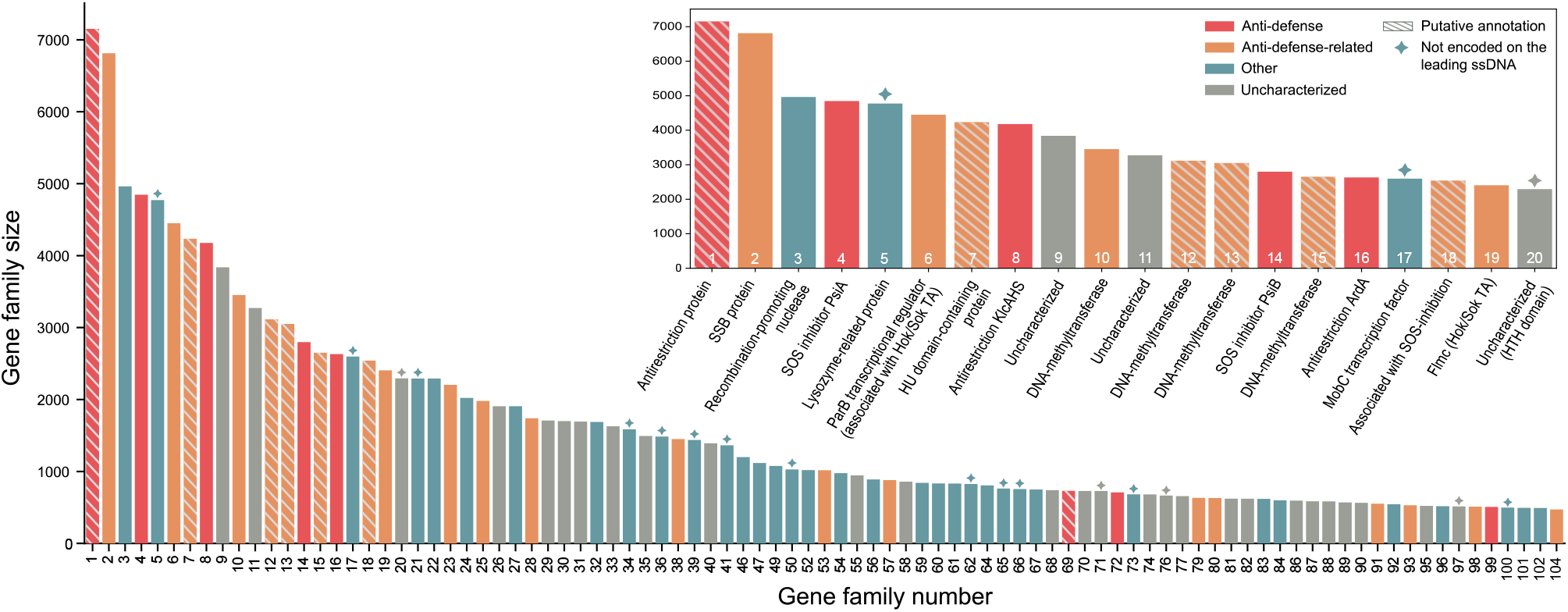
Gene families enriched in the leading regions. The largest gene families significantly enriched in the leading regions of conjugative elements were divided into four categories: 1) Anti-defense genes: anti-CRISPRs, anti-restriction genes, and SOS-inhibitors; 2) Anti-defense-related genes: methyltransferases, SSB proteins, toxins, and antitoxins; 3) Other: annotated genes without known association with anti-defense functions; 4) Uncharacterized genes. Colored diamonds (✦) indicate gene families encoded in the opposite orientation relative to the *oriT*, i.e., which cannot be transcribed from the leading ssDNA. Inset: Focus on the 20 largest gene families and their annotation. Putative annotations are indicated with striped bars.

We also frequently observed ssDNA-binding proteins (SSB) encoded in these regions, in most cases adjacent to SOS inhibitors (PsiA and PsiB). Several studies demonstrated the importance of plasmid-encoded SSB proteins for proper SOS-inhibition by PsiB^52,53^, and their possible role in plasmids evasion from the host CRISPR-Cas systems (through involvement in repairing double-strand DNA breaks)^54^. SSB proteins are also known to protect ssDNA intermediates from nuclease degradation. They directly interact with many different bacterial genome maintenance proteins, including recombination, repair, and replication proteins, such as polymerases^55–58^. These activities suggest they might have multiple protective functions in early conjugation stages.

Toxin and antitoxins were also highly represented in the leading regions, including both toxin-antitoxin (TA) systems and “orphan” antitoxins. Although TA systems are also encoded in other regions of conjugative element genomes, their overrepresentation in the leading region suggests a potential protective role in establishing plasmids and ICEs. TA systems could serve either as an “addiction system” of the conjugative element, a defense system against other MGEs^59–63^, or as an anti-defense system with antitoxins countering the function of host toxin-antitoxins, as previously demonstrated in bacteriophages^64,65^.

Notably, a major portion (36.26%) of the prevalent gene families in the leading region were uncharacterized. Given the considerable overrepresentation of anti-defense genes in this region, we expect that many of these uncharacterized families are likely to have anti-defense-related functions. Investigating the families lacking a clear functional annotation revealed numerous putative anti-defense-related functions, including anti-restriction activity, protection against nucleases, DNA repair, and association with the SOS-inhibition mechanism (see Table 1, Supplementary Text, and Supplementary Dataset 1).

**Table 1.**
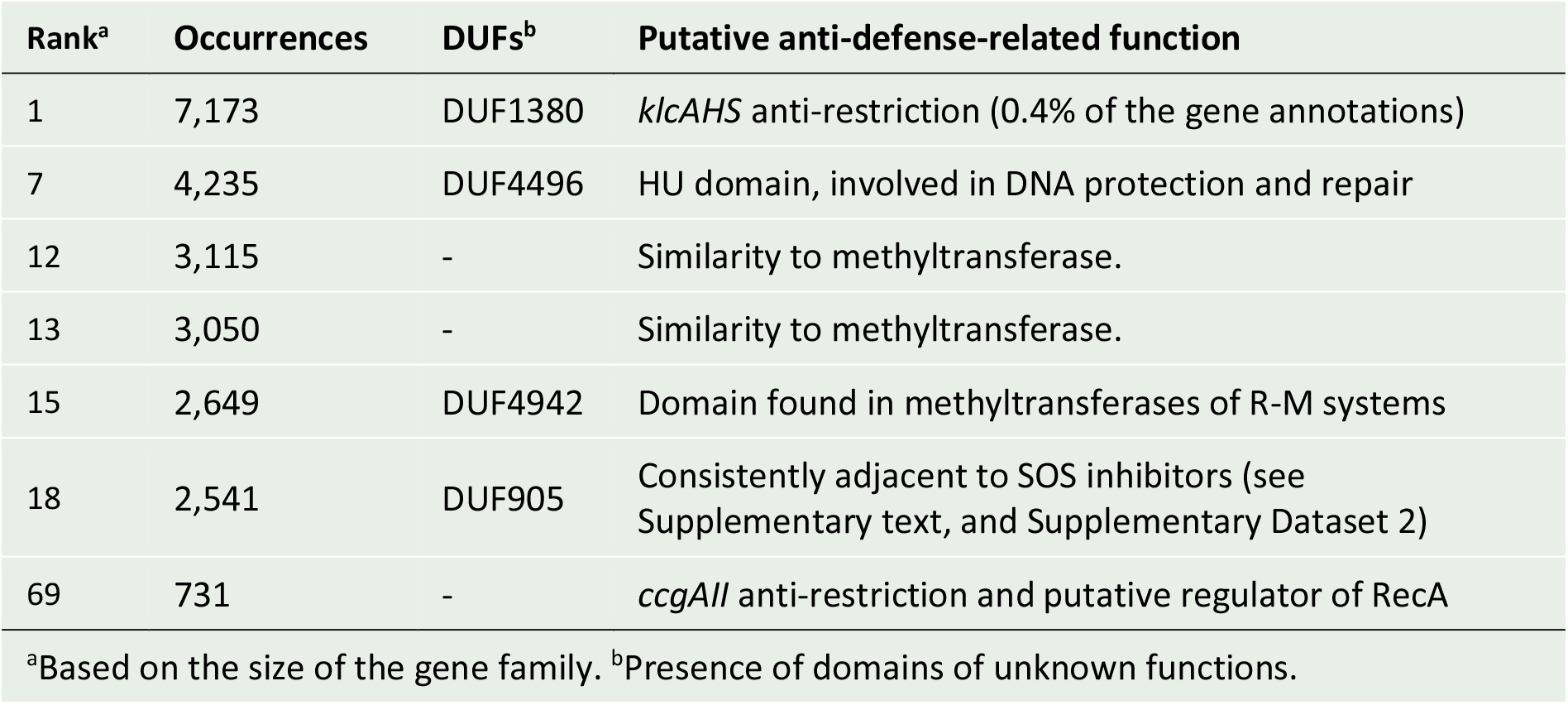
Uncharacterized gene families enriched in the leading region with putative antidefense functions.

### Anti-defense islands in the leading regions of conjugative elements

Examining the leading regions of conjugative elements revealed that the anti-defense genes tend to cluster into islands, as was reported for MGEs that contain clustered anti-defense genes^51^. We refer to these gene clusters as islands since most of the annotated genes in this region share a similar function, and they reside between defined boundaries: the *oriT* on one end and often *umuCD* homologs on the other. These anti-defense islands include different combinations of anti-defense and anti-defense-related genes adjacent to each other. For example, we located such an island in the leading region of a conjugative element of *Salmonella enterica* that contains two anti-CRISPR genes (*acrIC6* and *acrIF16*) in close proximity to gene encoding for anti-restriction (*klcAHS*), SOS-inhibitors (*psiA* and *psiB*), methyltransferases, ssDNA-binding proteins, and a TA system (*higB-higA*, Fig. 4). A similar island, with a few differences, was identified in the leading region of a plasmid from *Serratia marcescens*, an opportunistic pathogen, causing urinary tract, respiratory tract, and wound infections^66^. This island harbored an anti-CRISPR gene inhibiting a different type of CRISPR-Cas system (*acrIE9*) and an additional antitoxin gene (*hipB*, Fig. 4). The *hipB* antitoxin gene is usually located as part of *hipBA* operons and counters the toxicity of HipA^67^. However, in this anti-defense island, it was found next to a *higA/relE* TA system and may function as an “orphan” antitoxin inhibiting host TA defense systems.

**Figure 4.**
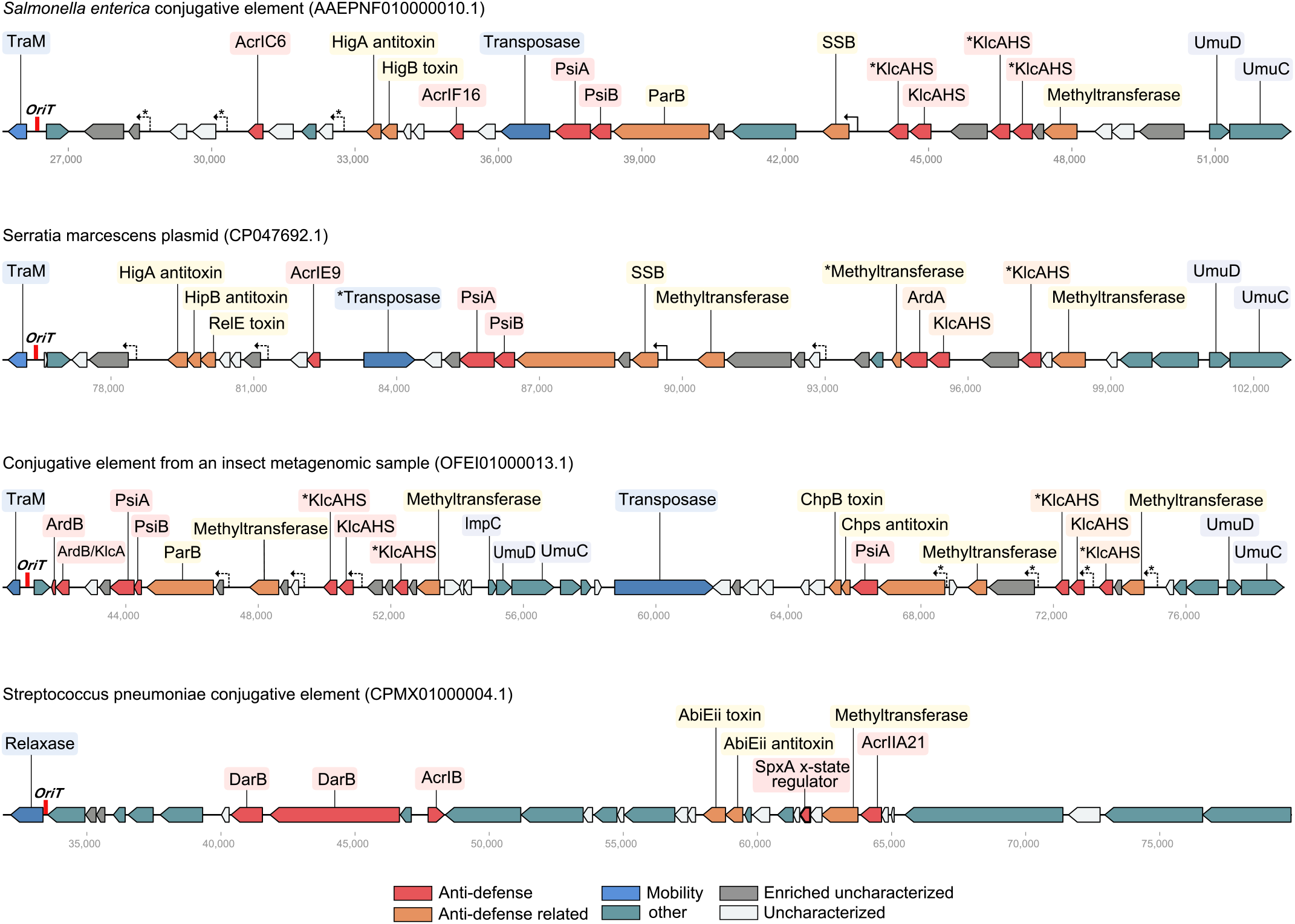
Representative anti-defense islands. Four representative anti-defense islands in leading regions of conjugative elements (additional islands presented in Supplementary Fig. 2). The *oriT* location is marked in red on the left. Genes are colored-coded according to their functional category: red: anti-defense, orange: anti-defense-related, blue: mobility (transfer genes), teal: gene without known association to anti-defense, grey: uncharacterized genes that were enriched in the leading regions, and white: other uncharacterized genes. F*rpo*-type promoters are indicated by an arrow. Promoters with significant similarity to known F*rpo* sequences are marked with a solid arrow, F*rpo* candidates that we identified are marked with a dashed arrow or dashed arrow with a star (*) for lower certainty candidates.

Curiously, many of the anti-defense islands were flanked by an operon of *umu*-like genes (also known as *mucAB*^68^ on plasmids) in the *oriT* distal region, which essentially forms the terminating boundary of the island (Supplementary Fig. 3). These genes are plasmids-encoded homologs of *umuC* and *umuD*, which form chromosomal translesion DNA synthesis polymerases (DNA polymerase V)^69^. Their role in plasmids is not yet clear^68^. While they are highly abundant in the leading region, they are not encoded in the orientation that would allow their expression from the ssDNA first transferred to the recipient bacteria (in 99.6% of the islands). Therefore, they are not expected to be expressed early upon conjugation. We noticed one case of a large anti-defense island (from a conjugative element recovered insect gut metagenomics) that appeared to consist of two adjacent islands separated by a transposase. An operon of the *umu*-like genes flanks each of these two adjacent islands (Fig. 4).

We wondered whether there were clusters of anti-defense genes that consisted primarily of uncharacterized gene families that we had identified as enriched in the leading regions. Supplementing our search with these uncharacterized gene families led to the detection of several additional anti-defense islands that could not have been detected based on our initial dataset of known anti-defense systems. One of the interesting islands we found owing to the enriched uncharacterized gene enriched was found in a conjugative element from *Streptococcus pneumoniae*. These bacteria can spread a pneumococcal disease in immunocompromised individuals and may cause hearing loss, brain damage, and death^70^. This *S. pneumoniae* conjugative element also harbors antibiotic-resistance genes against several antimicrobials, including tetracycline and chloramphenicol. The anti-defense island in the leading region of this element included a unique combination of anti-defense genes: a methyltransferase, two infrequent anti-CRISPRs (*acrIB* and *acrII21*), two anti-restriction genes (*darB*), a TA system (*abiEii-abiEi*), and two uncharacterized gene families prevalent in leading regions, close to a *spxA* gene (Fig. 4). SpxA is a transcriptional regulator involved in repressing the X-state, a general stress response mechanism in *S. pneumoniae* (a species lacking classical SOS-response pathway)^71^. As part of the stress response, competence genes are induced, and it has been shown that MGEs integrate specifically to these genes, disrupting transformation^72,73^. This is hypothesized to be a protective mechanism adopted by MGEs to prevent their elimination following the uptake of additional exogenous DNA^74^. The plasmid-encoded SpxA may thus serve as an “anti-X-state” protein that prevents the stress response, reminiscent of anti-SOS mechanisms in plasmids.

### Single-stranded DNA promoters are widespread in the anti-defense islands

Examining the orientation of the 91 gene families enriched in the leading region, we found that all the anti-defense and anti-defense related functions, without exceptions, were coded on the same strand relative to the *oriT* (Fig. 3). Specifically, the leading region anti-defense genes were consistently found on the anti-sense strand, such that they could be transcribed from the single strand that is first transferred to the recipient, even before the plasmid’s complementary strand is synthesized.

Transcription from ssDNA can be achieved using specific promoters, known as F*rpo* or *ssi* sequences, that create secondary DNA structures mimicking dsDNA, which enable recognition by RNA polymerase^15,21^. We thus sought known F*rpo*/*ssi* sequences in the leading regions of the 18,489 potential conjugative elements. We detected 11,840 F*rpo*-type homologous promoters in 5,341 conjugative elements. In each of the leading regions of the *S. marcescens* and the *S. enterica* plasmids, we identified one F*rpo-*type homologous sequence immediately upstream to the gene encoding the SSB protein, which is followed by SOS-inhibition genes (Fig. 5). Notably, the transcription by F*rpo* promoters is highly stimulated by SSB^20^.

**Figure 5.**
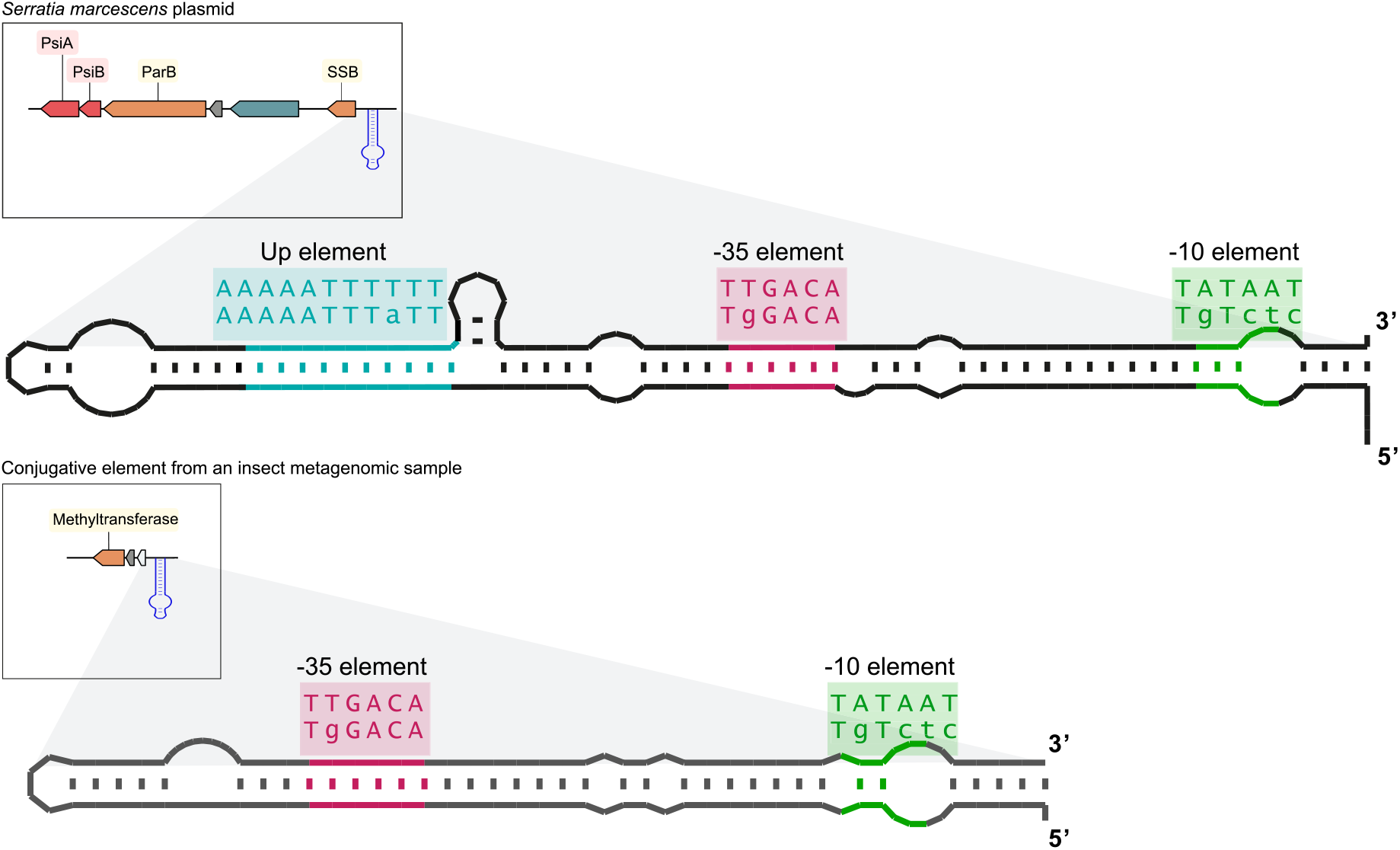
Single-stranded DNA promoters in the anti-defense islands. Top: Predicted secondary structure of the single-stranded F*rpo-*type promoter in *Serratia marcescens* plasmid. Bottom: A putative F*rpo* candidate in conjugative element recovered from an insect metagenome. The regions corresponding to −10, −35, and UP elements and complementary regions are colored. The sequence of these regions is highlighted above the structure, along with the canonical sequences of these elements. Upper case letters indicate nucleotides that conform with the canonical sequences of the −35 and −10 elements (5’-TTGACA-3’ and 5’-TATAAT-3’, respectively, for full sequences, see Supplementary Fig. 4).

In the two other islands depicted in Figure 4, no sequences homologous to F*rpo* were found. We postulated that additional ssDNA promoters might be present in the leading region to allow early expression of island genes. We searched for novel F*rpo*-type sequences in other regions upstream of ORFs in these islands. F*rpo-*type candidates were detected based on their predicted secondary structure and their conformance with known F*rpo* sequences and consensus sequences of the −35 and −10 elements (5’-TTGACA-3’ and 5’-TATAAT-3’, respectively; Supplementary Dataset 3). In the *S. marcescens* anti-defense island, we detected three putative F*rpo-*type candidates (F*rpo*’). In the *S. enterica* island, we identified three sequences bearing distant similarity to F*rpo-*type sequences (F*rpo*\*), with secondary structures similar to known F*rpo* sequences but considerable differences in the conserved −35 and −10 elements. Analyzing the islands from insect metagenome (Fig. 5) led to the detection of three additional Frpo-type candidates (F*rpo*’) and four putative F*rpo* candidates with only distant similarity (Frpo*) to known F*rpo* sequences.

We next searched for the putative F*rpo*-type candidates we found in the above-mentioned islands within the leading region of all 18,489 conjugative elements. We identified 5,768 additional F*rpo*’ and 736 F*rpo*\* candidates, presenting high and limited similarity to F*rpo* sequences correspondingly. Overall, the analysis of the regions upstream to ORFs in the islands revealed new and putative F*rpo*-type promoters in the leading regions of numerous conjugative elements. These results indicate that F*rpo* promoters are widespread in anti-defense islands and can potentially allow early expression from ssDNA during the very early stages of conjugation as part of the establishment phase to evade the host defense systems.

## Discussion

We found that the leading regions of plasmids and other conjugative elements are highly enriched with anti-defense genes. Examination of these regions across an extensive dataset of genomes and metagenomes revealed that various anti-defense systems in the leading region cluster together in “anti-defense islands”. These islands contain anti-CRISPRs, anti-restriction genes, and SOS inhibitors, alongside anti-defense-related genes, such as “orphan” methyltransferases, antitoxins, and SSB proteins. Uncovering anti-defense systems encoded in the leading regions suggests that these systems might be expressed very early upon entry to the recipient cell, well before the transfer is complete. This is supported by the presence of F*rpo*-type promoters, which allow expression from ssDNA. Early expression of anti-defense genes could provide rapid protection against the host defense systems during the initial establishment of the conjugative element in the recipient cell (Fig. 6). Currently, the genetic region adjacent to the propagation (mobility genes) and the *oriT* is designated as the “stability” or “establishment” region and includes numerous uncharacterized genes^75,76^. Our results indicate that inhibiting host defenses is a key function of genes in this region, and we thus propose designating it as “establishment and anti-defense” (Fig. 6).

**Figure 6.**
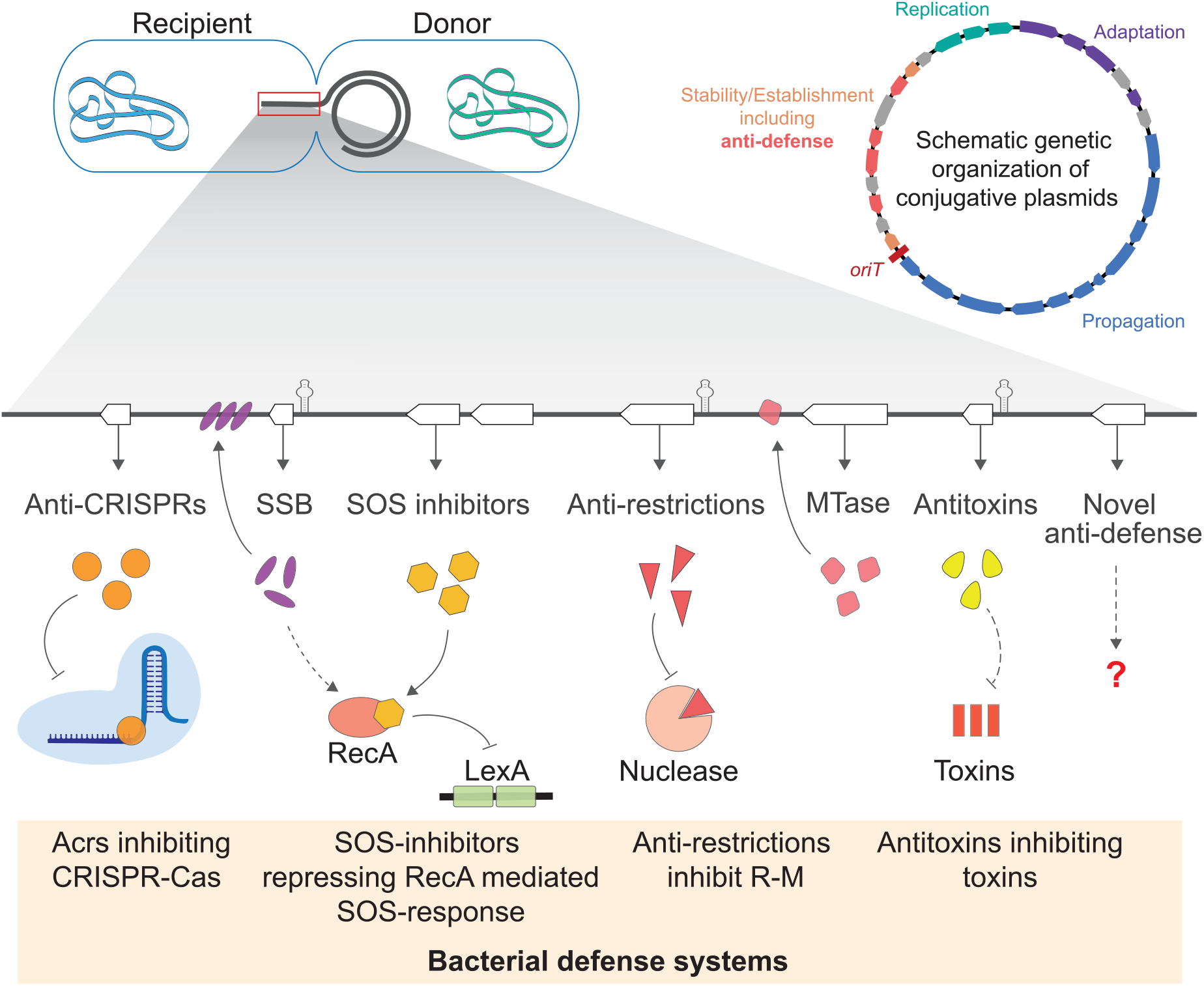
Suggested model of the protection provided to plasmids by diverse anti-defense genes encoded in the leading region. Owing to ssDNA promoters, the anti-defense genes can be expressed at the very early stages of transfer to a recipient cell. During this phase, the bacterial immune response recruits its defense systems to prevent the entry of the transferred foreign DNA. Anti-CRISPRs encoded on the plasmid can inhibit CRISPR-Cas systems; SOS-inhibitors, such as PsiB protein, can repress the cell SOS-response by preventing the activation of RecA thus inhibiting the cleavage of LexA, an SOS response transcriptional repressor; Single-stranded binding (SSB) protein are known to be involved in the SOS response inhibition mechanism, and may protect the transferred ssDNA from host nucleases. Methyltransferases (MTase), methylating the ssDNA can prevent recognition by the host restriction-modification (R-M) systems; Anti-restriction proteins can prevent DNA cleavage by R-M systems; and Antitoxins can neutralize host TA systems. In the top-right panel, a schematic genetic organization of a conjugative plasmid. The four main functional gene groups are colored: propagation (blue), adaptation (purple), replication (green), and the anti-defense genes (red) within the stability/establishment region (orange).

Numerous gene families enriched in the anti-defense islands were uncharacterized. However, their location in this region strongly indicates that they likely encode for anti-defense-related functions. Focusing the search on the leading region of conjugative elements could facilitate the discovery of new anti-defense systems, such as anti-CRISPR proteins, which are challenging to detect due to their small size and high variability^77,78^.

Intriguingly, a considerable fraction of the anti-defense islands are flanked by *umu*-like homologs, which encode for translesion DNA synthesis polymerases (Supplementary Fig. 3). These genes are widespread on conjugative elements^79–81^ and are also found in other MGEs, including the conjugative transposon Tn5252^82^, phages, and prophages^83,84^. Interestingly, they have been suggested to be involved in protecting plasmid and phage genomes by allowing DNA synthesis across otherwise unrepairable lesions^68,84^. In our analysis, they were almost exclusively encoded by the strand complementary to the leading region and are thus not expected to be expressed early with the anti-defense genes. Still, their abundance on various MGEs and their specific position at the edge of anti-defense islands in conjugative elements suggest they have a yet undetermined function in conjugation.

Our analyses were restricted to conjugative elements with a relaxase and a known *oriT* sequence, which were required to determine the transfer directionality. However, a large portion of the relaxase-containing sequences lacked a known *oriT* sequence, hampering our ability to comprehensively study potential conjugative elements^8,12^. Further, numerous small plasmids do not encode for relaxases and undergo transfer by exploiting the relaxosome of other conjugative elements *in trans*^85,6^. The robustness of the anti-defense gene location and F*rpo* promoters demonstrated in this study could be used to develop new approaches to identify previously unknown *oriT* sequences based on the location of the relaxase and the anti-defense islands. Similarly, the combination of *oriT* and anti-defense islands can be used to determine the leading region in small plasmids lacking a relaxase gene. These will allow studying additional plasmids and specifically investigating early expressed genes in small plasmids encoding only an *oriT*. Such small plasmids are important to study, as they have been shown to carry numerous antimicrobial resistance genes (ARGs) and make up at least half of all plasmids^5,8^.

Interestingly, several early expressed genes in phages have also been associated with anti-defense functions^35–39^. Given the complexity and variability of the anti-defense islands we have found in conjugative elements, it will be intriguing to examine genes that are early expressed both in phages and conjugative elements and explore their potential role in anti-defense.

An intrinsic part of the arms race between conjugative elements and their host is the interplay between bacterial defense systems and the anti-defense systems encoded on MGEs. We present a broad set of plasmid-encoded genes with anti-defense functions, supporting the hypothesis that anti-defense mechanisms among MGEs are diverse and dynamic, reflecting the large repertoire of bacterial immune systems. Our results provide a better understanding of strategies conjugative elements have developed during evolution to cope with the selective pressure of bacterial defense systems in a way that allows MGEs to rapidly overcome immunity and expand their distribution in the bacterial community^63,86^.

Notably, plasmids have been tested as potential conjugative delivery systems for various biotechnological applications, such as targeting antibiotic-resistance bacteria using CRISPR nucleases. However, these attempts resulted in low conjugation efficiency, especially in complex microbial communities like the human gut^87–89^. These studies emphasize that improving conjugation efficiency is vital for future applications. Our findings may provide a key factor to understanding the set of genetic tools required for efficient conjugation-based delivery systems for medical and biotechnological applications.

## Materials and Methods

### Datasets and initial annotation

The assemblies of all genomes and metagenomes from NCBI whole-genome projects (WGS)^41^ and all assembled metagenomes available from EBI Mgnify were downloaded on March 14, 2020^42^. After excluding genomes from Metazoa, Fungi, and Viridiplantae, the dataset included 596,338 genomes and 22,923 metagenomes from various ecosystems. This dataset contained more than 783 billion contigs of at least 10 kbp. In WGS, 31,119 sequences were explicitly annotated as plasmids. Gene calling and initial annotation were performed using prodigal^90^ version 3.0.0 and Prokka^91^ version 1.14.6.

### Relaxase/*traM* and *oriT* detection

Detection of relaxase and *traM* relaxosome genes was done using hmmsearch (HMMer^92^ version 3.3.2, e-value cutoff 1.00E-06) against all the proteins encoded by genomic and metagenomic sequences in our dataset. The profile HMMs were acquired from Pfam^93^ and MOBscan^94^ databases (Supplementary Dataset 4). Contigs with more than two relaxase or TraM hits were filtered out. Known *oriT* sequences were retrieved from oriTfinder^95^ (343 *oriT* sequences) and OriT-strast^96^ (112 sequences). The search for *oriT* sequences was performed using BLAST (BLAST+ 2.10.0, e-value cutoff 1.00E-06)^97^ against relaxase/*traM*-containing contigs (11,908 WGS plasmids and 1,019,093 genomes and metagenomes). Known *oriT* sequences were detected in 3,753 annotated plasmids with relaxase and in 196,414 relaxase-containing genomes and metagenomes. For contigs with more than one *oriT* sequence hit, we considered only the *oriT* with the best BLAST score. The distance between the relaxase/*traM* gene and the *oriT* was calculated as the number of nucleotides between the end of the relaxase/*traM* gene and the start of the *oriT*. We sought *oriT* that were in close proximity to the relaxase/*traM* gene. Thus, contigs in which this distance between the two was more than 3,500 bp were filtered out. We included contigs in which the *oriT* was partially contained within the relaxase gene (resulting in a negative distance), but cases in which the *oriT* was entirely contained within the relaxase gene were excluded. We also filtered out contigs in which relaxase genes or the *oriT* were at the ends of the contigs (first or last annotated sequences) since it impeded our ability to determine the relative location of the *oriT* and the relaxase/*traM* gene. Overall, this filtering process yielded 3,192 WGS annotated plasmids and 180,866 potential conjugative elements containing a relaxase/*traM* gene and an *oriT*.

### De-duplication of redundant sequences

To avoid duplicate sequences, we clustered all 491,157 ORFs of the 3,192 WGS plasmid contigs containing relaxase and *oriT* using CD-HIT^98^ (version 4.6). According to this clustering, we calculated the percentage of shared ORFs for each pair of contigs. If two plasmids shared more than 90% of the ORFs, the plasmid with fewer ORFs was filtered out. This process yielded 1,554 representative plasmids. We performed the same de-duplication process for 180,866 potential conjugative elements we identified in genomic and metagenomes sequences, yielding 17,151 non-redundant contigs of potential conjugative elements. Combining the plasmids with the rest of the potential conjugative elements resulted in a total of 18,489 non-redundant contigs of potential conjugative elements. The host phylogenetic distribution of these non-redundant contigs (Fig. 2d) was mapped to the bacterial subtree of iTol^47^ and generated using ggtreeExtra^99^.

### Anti-defense and mobility gene annotation

Protein families with known anti-defense functions were modeled using 120 profile HMMs (Supplementary Dataset 4). To characterize the plasmid’s transfer genes, we searched for specific conjugation proteins, such as type IV secretion system proteins, using HMMs downloaded from Pfam^93^ or computed based on proteins from relevant KEGG orthologs^100^ (Supplementary Dataset 4). To identify transposases, we used 49 HMMs from TnpPred data archive^101^. Hmmsearch (with an e-value cutoff of 1.00E-06) was performed against all non-redundant potential conjugative elements sequences containing a relaxase/*traM* and an *oriT*.

### Statistical analysis and ORF clustering

To test which ORF positions in the leading region of plasmids were significantly enriched with anti-defense genes, we performed a Fisher’s exact test (one-sided, p-value < 0.05) on the anti-defense gene count at each location summing over a sliding window of five ORFs (anti-defense gene count versus the total number of genes). The same test was performed separately for each anti-defense category (namely anti-CRISPRs, anti-restriction genes, and SOS inhibitors). This analysis was performed on the 1,554 sequences of annotated plasmids for positions with at least 50 ORFs. The p-values were corrected for multiple testing using FDR (alpha = 0.05).

To cluster the genes in the leading region (first 30 ORFs) of all non-redundant potential conjugative sequences, we used MMseqs2^102^ (with sensitivity of 0.75 and coverage of 0.5). The ORFs in 105 gene families with more than 450 ORFs were aligned using MAFFT^103^ (version 7.475), and an HMM was constructed from each alignment (Supplementary Dataset 5). Hmmsearch (e-value cutoff 1.00E-06) of these HMMs was performed against all potential conjugative sequences. To statistically test the enrichment of each gene family in the first 30 ORFs in the leading region, we performed a one-sided Fisher’s exact test and calculated the p-values after FDR correction (alpha = 0.001). Fourteen of the 105 gene families were found not to be significantly enriched in the leading regions and were omitted from downstream analyses.

We searched for known conserved domains within the hypothetical gene families using NCBI CDD^104^ and HHpred^105^ (databases: PDB^106^, Pfam^93^, TIGERFAMs^107^; e-value cutoff 1.00E-10). ORFs were annotated based on significant hits to conserved domains. In gene families with ORFs that received different annotations, the most frequent annotation in the gene family was used.

In each of the 91 significantly enriched gene families, we examined the orientation of each of its ORFs relative to the *oriT* position. The overall orientation of the gene family was defined based on the majority of its ORFs.

The log-odds co-occurrence ratio of each pair of gene families was calculated as the log (base 2) of the ratio of the frequency of co-occurring gene pairs to the product of the frequencies of each gene separately (Supplementary Dataset 2).

### F*rpo* and *ssi* promoter identification

To identify known *Frpo/ssi* sequences in our anti-defense islands, we created a BLAST^97^ dataset of all the intergenic regions larger than 350 bp in the leading regions of all potential conjugative elements. We then performed a BLAST search (BLAST+ 2.10.0, e-value cutoff 1.00E-06) against five known *Frpo/ssi* sequences^16,108^ (Supplementary Dataset 3).

New candidate F*rpo* sequences were detected by seeking the consensus sequences of the −35, −10 (5’-TTGACA-3’ and 5’-TATAAT-3’, respectively), and the UP-element regions in the intergenic regions of the islands represented in Fig. 4 (Supplementary Dataset 3). We then performed a BLAST search of the putative F*rpo* candidates found in these islands against all the leading regions of all potential conjugative elements.

The DNA secondary structures of the F*rpo*/*ssi* elements were calculated using the RNAfold web server^109,110^ with the 2004 David H. Mathews model for DNA. The graphical illustrations of the RNA structures (Fig. 5) were produced using RNAtist^111^.

## Acknowledgments

We wish to thank Dr. Karin Mittelman, Prof. Gil Segal, Prof. Eliora Ron, Prof. Avigdor Eldar, Prof. Adi Stern, and Prof. Uri Gophna for their helpful discussions and comments on the manuscript. We are thankful to Danielle Miller for her assistance in the bioinformatic analysis. This research was partly funded by ISF grant number 1692/18.

## Author contribution

B.S. and D.B. conceived and designed the study, performed the data analysis, and wrote the manuscript.

