## Supplementary Material for "A diverse repertoire of anti-defense systems is encoded in the leading region of plasmids"

### Supplementary Text

Putative anti-defense-related functions of uncharacterized genes enriched in the leading region

The following uncharacterized gene families were enriched in the leading region of conjugative elements and were found to have potential anti-defense-related functions (see also Table 1 and Supplementary Dataset 1):

**Family #1 (7,153 genes):** The most prevalent family enriched in the leading region consisted mostly of uncharacterized genes. However, a minute fraction of the genes in this family (0.4%) was annotated as *klcAHS* anti-restriction genes, suggesting this family might be a yet uncharacterized anti-restriction gene family.

**Family #7 (4,235 genes):** This family includes DUF4496, also known as the "CCDC81 family HU domain 1", found in eukaryotes[1]. HU proteins are part of the nucleoid-associated proteins (NAPs), together with FIS, H-NS, and IHF proteins[2]. HU proteins can help prevent DNA damage and are also able to protect DNA against nucleases[3,4,5]. This suggests HU proteins may also be part of the anti-defense mechanisms of plasmids.

**Family #12 (3,115 genes):** This family seems to be a remote homolog of methyltransferases (HHsearch e-value $2\times10^{-35}$).

**Family #13 (3,050 genes):** This family seems to be a remote homolog of putative methyltransferases (HHsearch e-value $3\times10^{-58}$).

**Family #15 (2,650 genes):** This family contains the DUF4942, found in DNA methyltransferases in restriction-modification (R-M) systems, and was suggested to be a novel group of methyltransferase[6].

**Family #18 (2,541 genes):** This family encodes DUF905, also known as the *ykfF*, a gene found next to SOS inhibitors on pKSR100 plasmid[7] and several IncI plasmids[8]. Accordingly, we also found that, in 99.05% of their occurrences, genes of this family are located on the same island as the PsiA SOS inhibitor (gene family 4) and in 98.54% of the cases with SSB protein (gene family 2), which is known to play part in the SOS inhibition mechanism (see Supplementary Dataset 2).

**Family #69 (731 genes):** The *ccgAII* gene. The function of this gene is not entirely clear, but it was implicated in protecting from the type I restriction system[9,10,11] and shown to play a role in regulating RecA overproduction[12]. Both reported functions suggest it might represent another anti-defense mechanism.

### Supplementary Figures

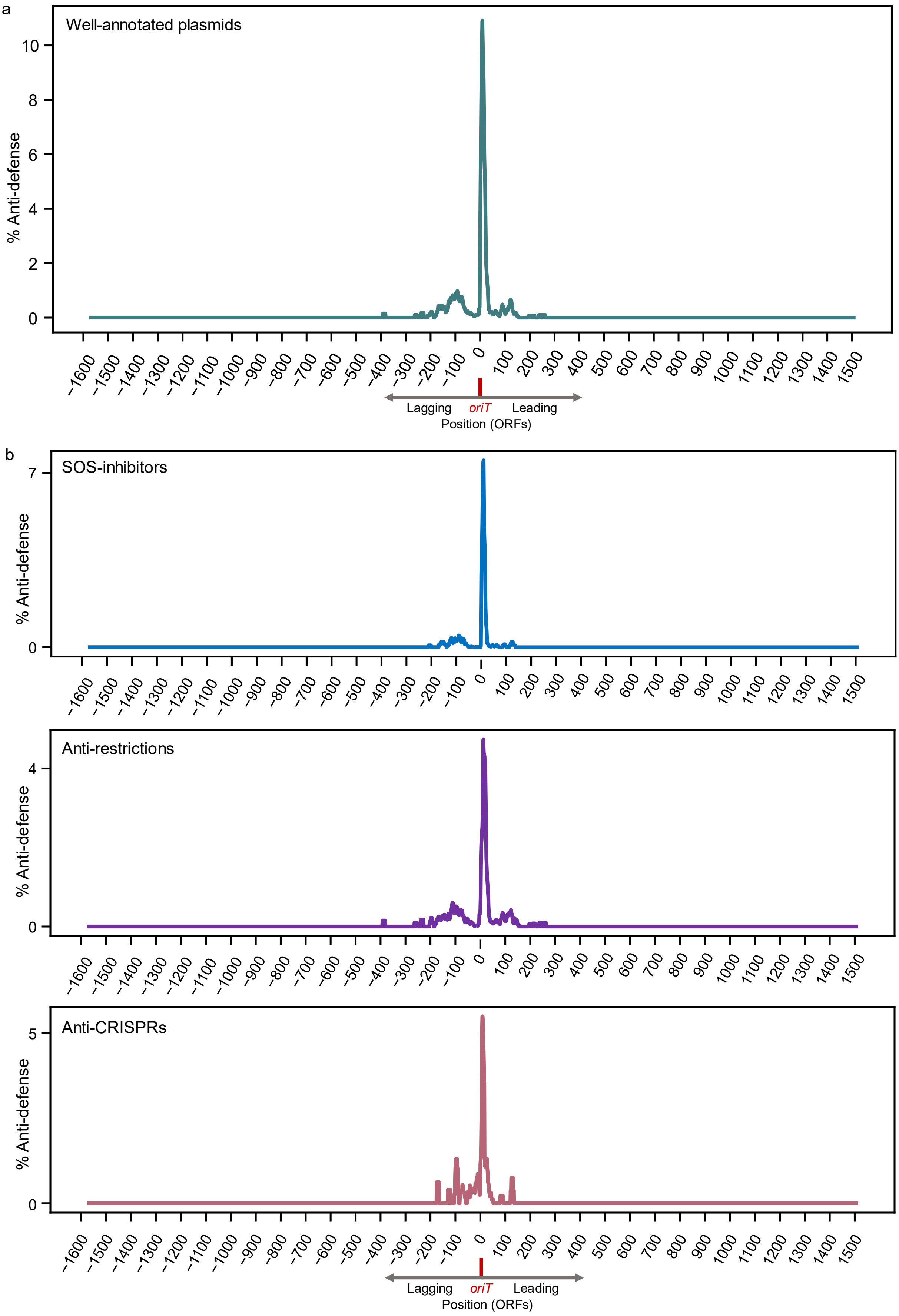

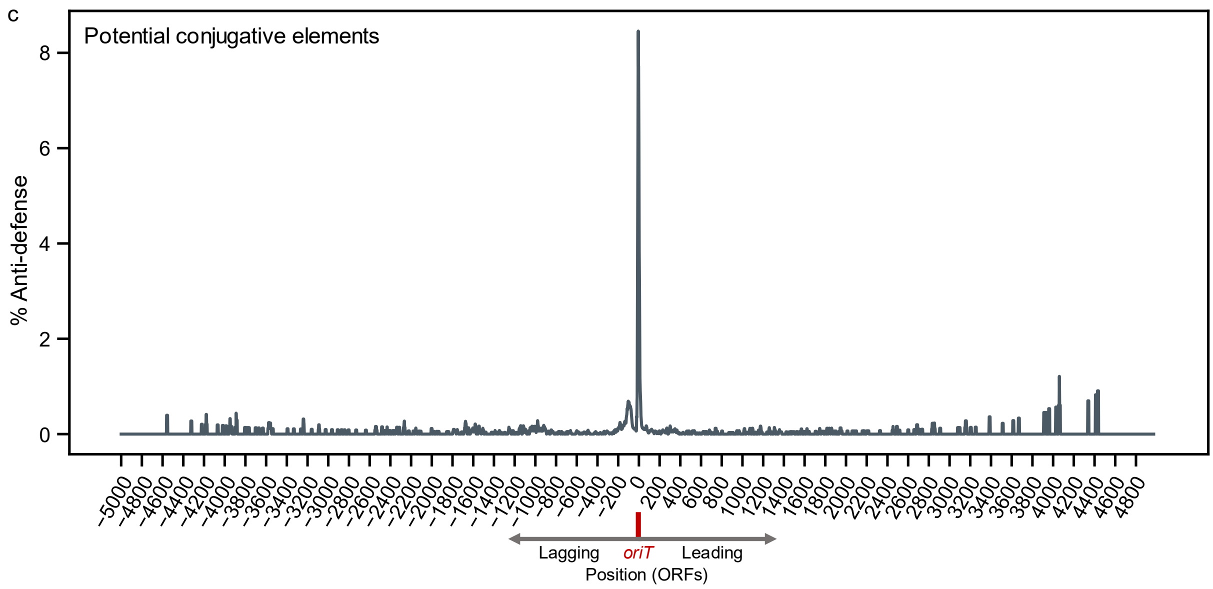


**Supplementary Figure 1 | Frequency of anti-defense genes relative to the origin of transfer (*oriT)*, full data. (a)** Analysis of anti-defense genes encoded on sequences explicitly annotated as plasmids. The ORF indexes on the *x*-axis are according to their position relative to the *OriT*, such that 0 is the first ORF in the leading region. The *y*-axis denotes the average frequency of anti-defense genes, combining well-characterized SOS inhibitor, anti-restriction, and anti-CRISPR genes over a window that includes five ORFs upstream and downstream. **(b)** Breakdown of the anti-defense gene frequency according to their functional categories: SOS inhibitors, anti-restrictions, and anti-CRISPRs. **(c)** Analysis of anti-defense gene frequency from potential conjugative elements retrieved from genomic and metagenomic databases.

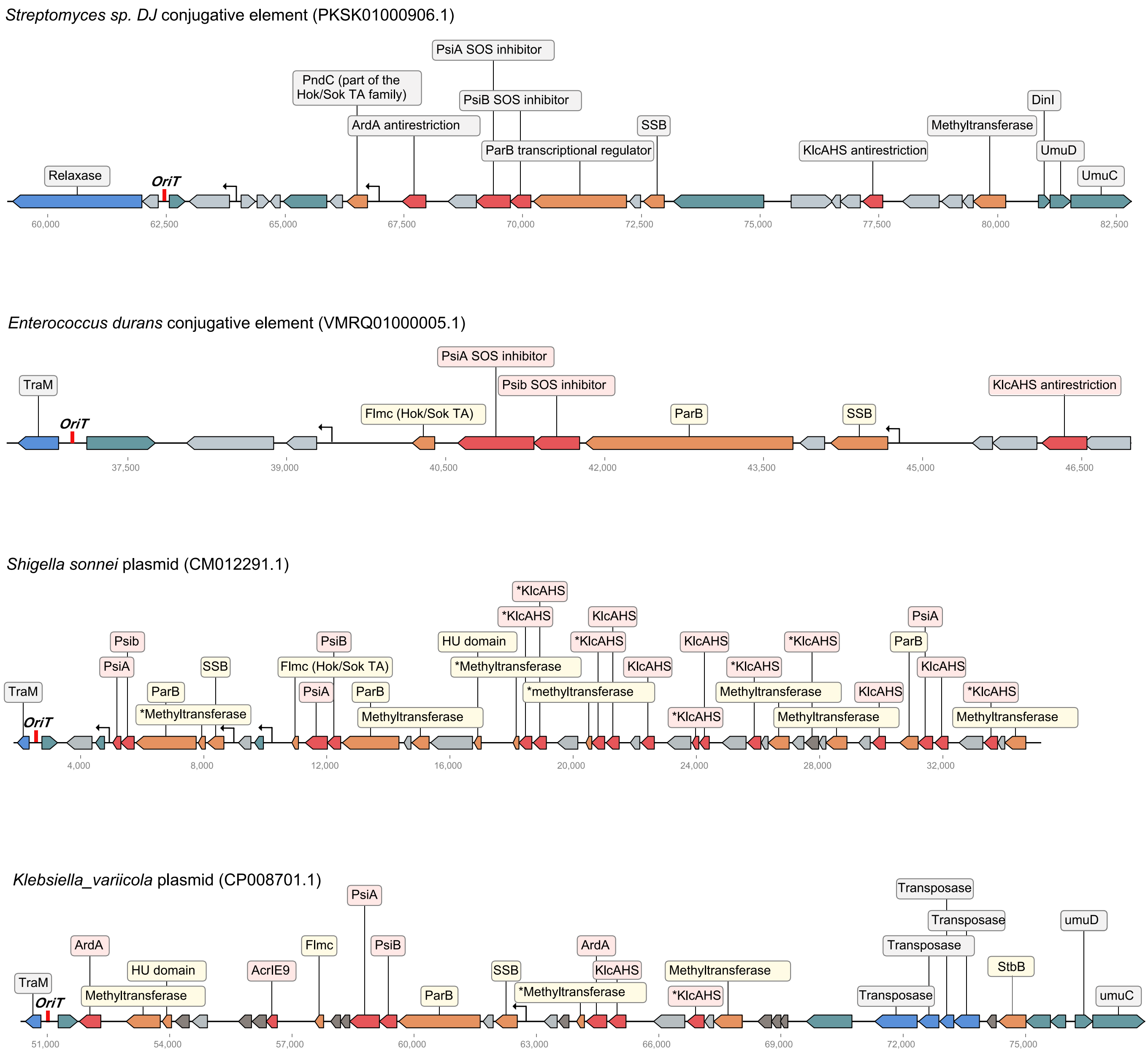


**Supplementary Figure 2 | Additional examples of anti-defense islands found in conjugative elements from various bacterial hosts.** The *oriT* location, where the conjugation transfer starts, is marked in red on the left. Genes are colored-coded according to their functional category: red: anti-defense, orange: anti-defense-related, blue: mobility (transfer genes), teal: gene without known association to anti-defense, grey: uncharacterized genes that were enriched in the leading regions, and white: uncharacterized genes. F*rpo*-type promoters are indicated by an arrow.

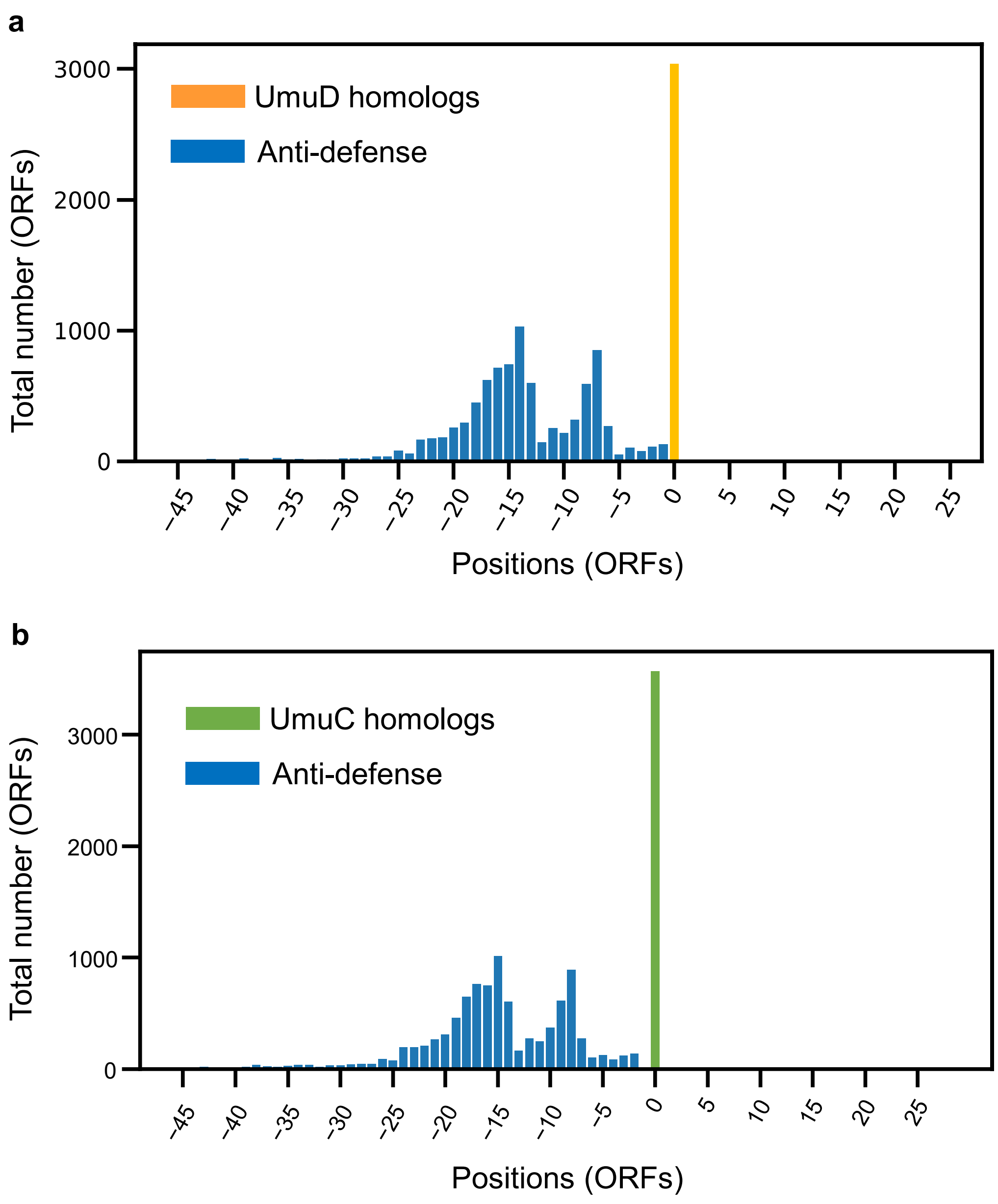


**Supplementary Figure 3 | Position of the anti-defense genes in the leading region of 18,489 potential conjugative elements relative to *umuCD* homologs.** (a) The position of *umuD* homologs was defined at position 0, and the total number of anti-defense genes at each relative position was calculated. **(b)** A similar analysis was for *umuC* homologs. In cases where there were several *umuD*/*umuC* genes in the same leading region, we considered the one closest to the *oriT*.

**a**

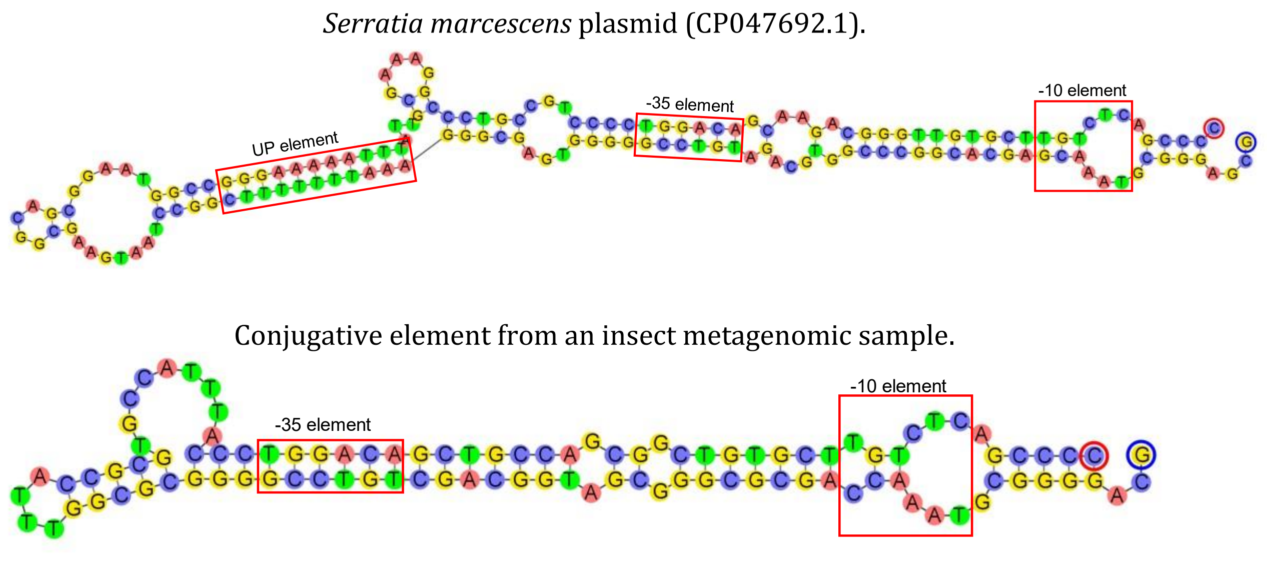


**b**

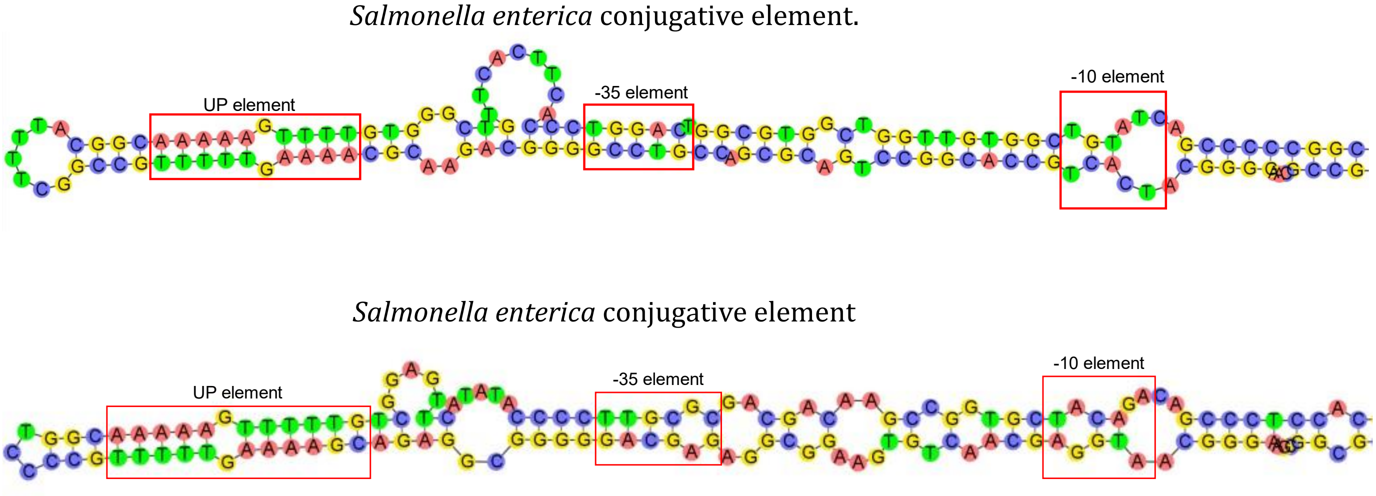


**Supplementary Figure 4 | Sequences and predicted secondary structures of F*rpo* promoters presented in Fig. 4b. (a)** F*rpo* found upstream to SSB protein in *S. marcescens* plasmid; and candidate F*rpo* we detected in conjugative element recovered from an insect gut metagenome. Both had high sequence similarity to known F*rpo* sequences **(b)** Candidates F*rpo* sequences, with limited similarity to known F*rpos*, that were found in *S. enterica* conjugative elements. The regions corresponding to -10, -35, and UP elements are indicated with a red square.
